# Quantitative profiling of millions of nucleotides reveals sequence-encoded interactions that govern plasmid propagation

**DOI:** 10.64898/2025.12.15.694402

**Authors:** Tom Copeman, John H.C. Fong, Joshua Mayne, Thomas E Gorochowski, S Rodriguez-Conde, Tom Ellis, Francesca Ceroni

## Abstract

Plasmids are central to modern biotechnology, especially therapeutic development, yet their propagation in *Escherichia coli* remains difficult to predict. Although expression-induced burden is well understood and can be mitigated, the impact of foreign DNA segments that do not function in bacteria on plasmid propagation and stability remains largely unknown. Here we developed a pooled, sequencing-based framework that performs quantitative profiling across millions of bases, enabling high-resolution assessment of plasmid fitness at scale and revealing cryptic, sequence-encoded interactions between foreign DNA elements and bacterial hosts. Promoter-like motifs, transcription factor binding site homology, and recombination-prone architectures emerge as major determinants of propagation efficiency, with context dependent effects demonstrating that plasmid behaviour arises from higher-order interactions between parts rather than isolated elements. Extending this framework, we introduce TRACE, a neural network model trained on degenerate sequence libraries that predicts plasmid propagation directly from sequence. TRACE generalises across plasmid architectures and can be fine-tuned on experimental datasets to improve predictions of manufacturability and host compatibility. These advances establish a generalisable, data-driven framework for understanding and designing host-aware plasmids, transforming plasmid production from an empirical process into a predictable property of DNA sequence.

## Introduction

Plasmids are the central genetic vehicles of biotechnology, enabling nearly every workflow from cloning and library expansion to heterologous gene expression across bacterial, yeast, and mammalian systems. Their versatility has made them indispensable not only in academic research but also across industrial biotechnology and therapeutic manufacturing, where plasmid DNA (pDNA) forms the molecular backbone for viral vector production, mRNA vaccines, and cell and gene therapies.^(1)^ Large-scale production of these plasmids still depends on *Escherichia coli* as the propagation host, valued for its genetic tractability, rapid growth, and well-established GMP infrastructure.(^2, 3^)

Plasmid maintenance imposes a measurable physiological cost on *E. coli*. Replication, transcription, and translation of plasmid-encoded genes divert the host’s energetic and biosynthetic resources, nucleotides, ribosomes, polymerases, and ATP, away from essential processes. This can trigger characteristic stress responses when the burden becomes excessive.^(4-7)^ Such impact has already been well characterised in synthetic biology, and many mitigation strategies have been proposed, including resource allocators that control ribosome availability,^(8)^ feedback loops that down-regulate heterologous gene expression, ^(9)^ and genome minimisation to reduce competing load. ^(10)^

As synthetic biology expands into mammalian systems, plasmid designs now routinely incorporate viral and eukaryotic genetic parts that must still be propagated in *E. coli*. Frameworks such as the Mammalian Tool Kit (MTK) exemplify this shift, enabling modular, multi-part assembly of mammalian expression cassettes for rapid prototyping.^(11)^ Although these parts are not designed to function in bacteria, they can nonetheless engage the host’s cellular machinery in unexpected ways (**Fig. 1A**). Recent evidence suggests that plasmid sequences can actively interfere with bacterial physiology through unintended molecular interactions. AT-rich eukaryotic motifs can resemble σ^70^ promoter elements and trigger spurious transcription, ^(12-18)^ viral or synthetic DNA can carry partial matches to bacterial transcription-factor binding sites that misdirect global regulators,(^19, 20^) and repetitive or homologous elements can recombine to form concatemers, compromising plasmid stability and segregation. ^(21-24)^ These interactions highlight that plasmids are not passive carriers of genetic information but dynamic participants in the regulatory, structural, and metabolic landscape of the host cell.

**Figure 1.**
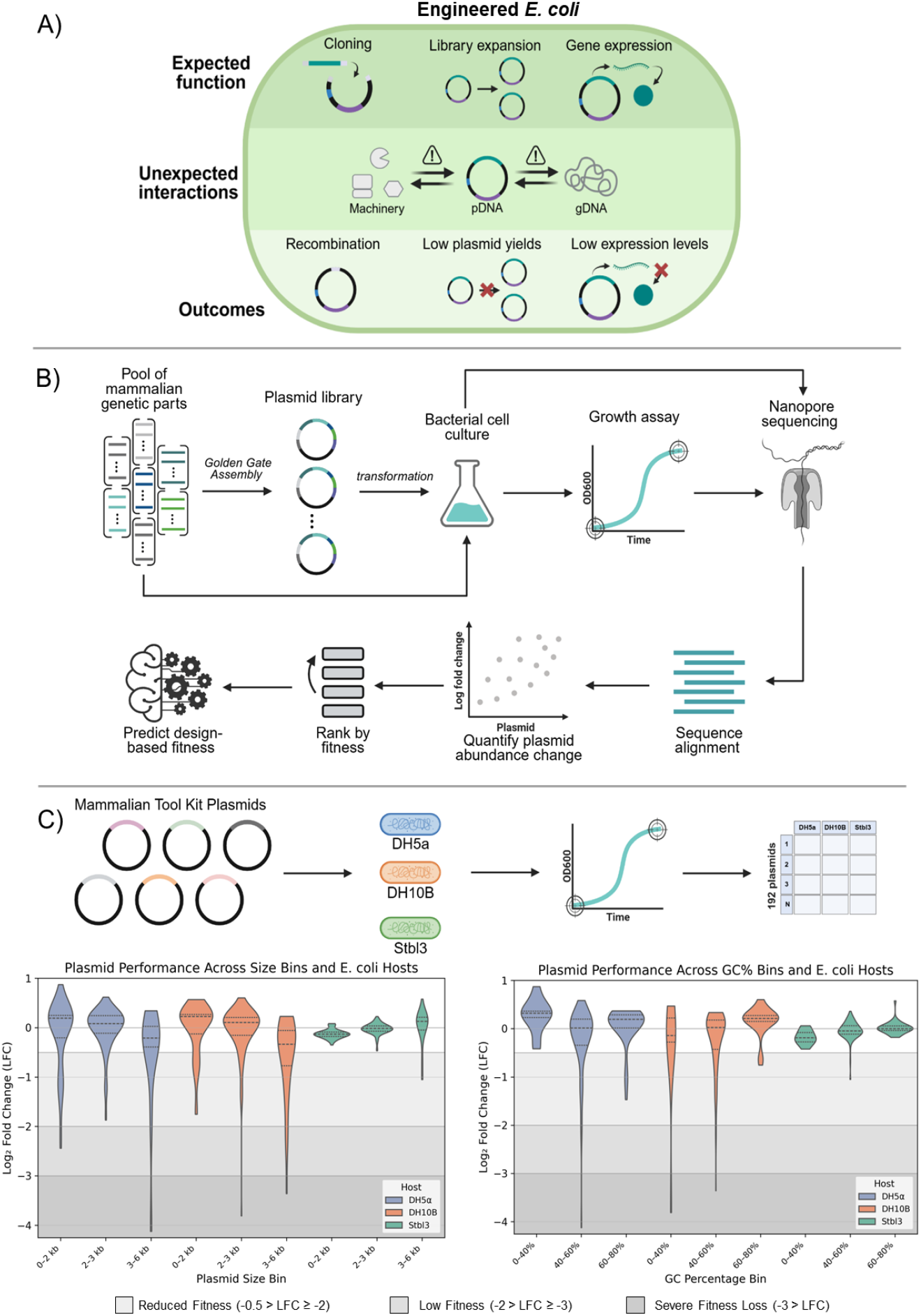
Systematic assessment of plasmid–host interactions and plasmid fitness across bacterial strains. A) Plasmids support cloning, library propagation, and gene expression, but unintended interactions with host transcriptional and replication machinery can drive recombination, plasmid instability, low plasmid DNA (pDNA) yields, and reduced expression. B) Overview of the pooled plasmid fitness assay. A diverse library of mammalian expression plasmid parts is assembled using Golden Gate cloning and transformed into *E. coli*. The pooled culture is grown over time, and changes in plasmid abundance are measured via long read nanopore sequencing and alignment. Log fold change (LFC) in relative plasmid abundance is used as a fitness metric and ranked across the library. Machine learning models predict sequence-dependent fitness effects. C) A set of Mammalian Toolkit (MTK) plasmids was plots show plasmid fitness with three distinct categories of poor plasmid propagation including reduced fitness (light grey), low fitness (grey) and severe plasmid loss. This was compared to plasmid size (left) and GC content (right).

Despite these insights, no systematic framework exists for predicting how specific DNA features influence plasmid maintainability, stability, or yield in *E. coli*. Most industrial and research workflows still rely on empirical design–test cycles, and plasmids that fail to amplify are often discarded without mechanistic explanation. The lack of predictive understanding hampers optimisation of high throughput prototyping pipelines and limits the scalability of pDNA manufacturing as plasmids become increasingly complex and further diverge from bacterial genomic norms.

Here, we address this gap by developing a pooled parallel assay that utilises next-generation sequencing to quantitatively profile plasmid fitness during competitive growth in *E. coli*. Using this framework, we measure the effects of millions of bases of DNA in *E. coli*, investigating how libraries of mammalian parts, and subsequently, combinatorial multi-part expression cassettes, propagate within a pooled cell culture. By integrating fitness measurements with transcriptional profiling, long-read structural analysis, and computational motif discovery, we map how distinct sequence features interact with the bacterial host machinery. Finally, by extending these principles to an unbiased library of fully degenerate sequences, we derive general rules governing plasmid compatibility in *E. coli*. The curated data trained a neural network that formed the basis of TRACE, a predictive platform for forecasting plasmid propagation directly from the DNA sequence alone. This establishes an experimentally grounded, host-aware framework to enable future rational plasmid design.

## Results

### Assay Development and Validation

To enable the high-throughput characterisation of plasmid–host interactions, we developed a pooled assay that measures the propagation efficiency of plasmid libraries, simultaneously. Plasmids were tested as they came in the MTK or were assembled by Golden Gate assembly before being transformed into *E. coli* and propagated in 50 mL batch culture under antibiotic selection for 16 hours (**Fig. 1B**). Plasmid DNA was harvested at the start (t = 0 h) and end (t = 16 h), sequenced using long-read nanopore sequencing, and plasmid fitness was quantified as the log fold change (LFC) in relative read abundance between these timepoints. This pooled format enabled direct comparison of maintainability across hundreds of constructs within the same culture environment, avoiding the throughput and variability limitations of isolated growth assays.

We first verified that Nanopore read counts provide an accurate measure of relative plasmid abundance. To do this, we prepared a panel of plasmids at known concentrations, quantified using the Qubit dsDNA High Sensitivity assay, and sequenced them together in a pooled run. Read count per plasmid correlated strongly with Qubit-measured plasmid concentration (**Fig. S1A**), confirming that Nanopore read abundance reflects actual plasmid abundance in a mixed pool. Next, to test whether plasmid length influenced read count independently of concentration, we prepared plasmids of different sizes (ranging from <2 kb to >10 kb) but normalised them to equal DNA concentration before sequencing. Read counts remained similar across plasmid sizes (**Fig. S1B**), demonstrating that Nanopore abundance estimates are not biased by plasmid length under our sequencing conditions.

To relate sequencing-derived abundance changes to established measures of plasmid burden, we compared plasmid LFC against standard physiological and molecular readouts. OD_600_ growth rate and maximum culture density are commonly used as proxies for global cellular burden and stress,^(6)^ while plasmid copy number (PCN) measured by qPCR is a widely used metric for assessing plasmid replication efficiency and stability.^(25)^ Accordingly, we monitored cell growth dynamics by OD_600_ over a time course assay and measured the resulting PCN by qPCR (**Fig. S2**). This allowed us to evaluate how sequencing-measured fitness relates to conventional measures of plasmid behaviour. Plasmid fitness correlated strongly with PCN (ρ = 0.71, p = 0.002) indicating that the LFC metric reflects the ability of plasmids to be stably replicated and maintained. In contrast, correlations with maximum culture density (ρ = 0.24, p = 0.37) and growth rate (ρ = 0.29, p = 0.27) were weak, suggesting that variation in plasmid fitness is not primarily driven by changes in bulk cell growth physiology. Instead, LFC captures plasmid-specific maintenance costs, such as replication stability and segregation efficiency, rather than indirect reductions in overall growth rate. This demonstrated that the pooled parallel assay provides a biologically grounded, high-throughput proxy for plasmid maintainability, consistent with conventional burden quantification approaches but scalable to hundreds of constructs simultaneously.

The pooled propagation assay first screened 192 plasmids taken from the MTK system, sourced from Addgene (see Methods section). Each plasmid contained a single mammalian regulatory, or coding sequence cloned into the same pUC-derived backbone, ensuring that differences in plasmid behaviour only reflected the properties of the insert. To assess the impact of host genotype, plasmid propagation was compared across three commonly used *E. coli* strains with differing recombination and stability phenotypes, DH5α, DH10B, and Stbl3.(^26-28^)

Across these strains, the majority of MTK parts exhibited neutral fitness, while a minority disrupted plasmid propagation. In DH5α and DH10B, 22 to 31 parts resulted in reduced plasmid fitness, 3–4 low fitness, and two severe losses, whereas Stbl3 was largely permissive with only four reduced fitness designs and no low or severe outcomes (**Fig. 1C**). This difference is consistent with the design of Stbl3 as a strain optimised for unstable or repetitive DNA, incorporating reduced recombination activity, lower insertion-sequence mobility, and routine propagation at reduced temperature, all of which act to stabilise plasmids in bacterial hosts. Even in this most permissive background, however, certain designs remained vulnerable to losses in plasmid fitness. (^28, 29^)

Plasmid size and GC content are known from prior work to influence cell burden, plasmid copy number and transcriptional behaviour in bacteria and were therefore examined as background trends in the dataset (**Fig. 1C**; **Fig. S3**).(^16-18, 30, 31^) Larger inserts were associated with poorer fitness in DH5α (ρ = −0.36, p < 0.001) and DH10B (ρ = −0.42, p < 0.001), whereas Stbl3 showed the opposite trend, with larger plasmids better tolerated (ρ = 0.57, p < 0.001), indicating that size influences plasmid behaviour in a strain-dependent manner rather than acting as a universal constraint. GC content likewise showed no consistent relationship with fitness across hosts, with no effect in DH5α but a weak positive association in DH10B (ρ = 0.32, p < 0.001) and Stbl3 (ρ = 0.35, p < 0.001). Importantly, neither size nor GC content separated the poorly performing plasmids from the remainder of the library. Low- and severe-fitness constructs were observed across all bins, with no enrichment confined to extreme size or GC ranges. This indicates that while broad sequence properties shift population-level fitness distributions, they do not explain the most severe failures.

Against this background of gradual size- and composition-related effects, four plasmids consistently performed poorly across all strains including MTK3_009 (PHYB-VP16), MTK3a_026 (dSaCas9-NLS), MTK3b_019 (dSaCas9), and MTK8b_004 (5′ HIV LTR), all of which contain parts associated with high regulatory activity or complex protein expression. This indicated that the shared features in these parts were responsible for the impaired plasmid stability and maintenance independent of host, size, or base composition, thus pointing to the possibility of unintended negative interactions between mammalian DNA and the *E. coli*. Although prior studies have reported isolated incompatibilities between viral or eukaryotic sequences and *E. coli*,(^13, 15^) our workflow highlights that such effects are widespread and measurable at scale. The pooled propagation assay therefore provides a robust framework for resolving plasmid–host interactions and for investigating how genetic context shapes plasmid behaviour.

Although plasmid size and GC content contributed to broad trends in fitness across the MTK library, they did not account for all constructs that propagated either unexpectedly poorly or unexpectedly well (**Fig. 1C**). This indicated that more specific sequence-encoded interactions between foreign DNA and the *E. coli* machinery were influencing plasmid propagation. To dissect these effects, we investigated three mechanistic routes previously investigated in heterologous pDNA in bacterial hosts, 1) unintended transcriptional activity driven by cryptic promoter elements, 2) structural instability through concatemer formation, and 3) sequestration of global transcription factors involved in host metabolic regulation (**Fig. 2A**).^(13-15)^

**Figure 2.**
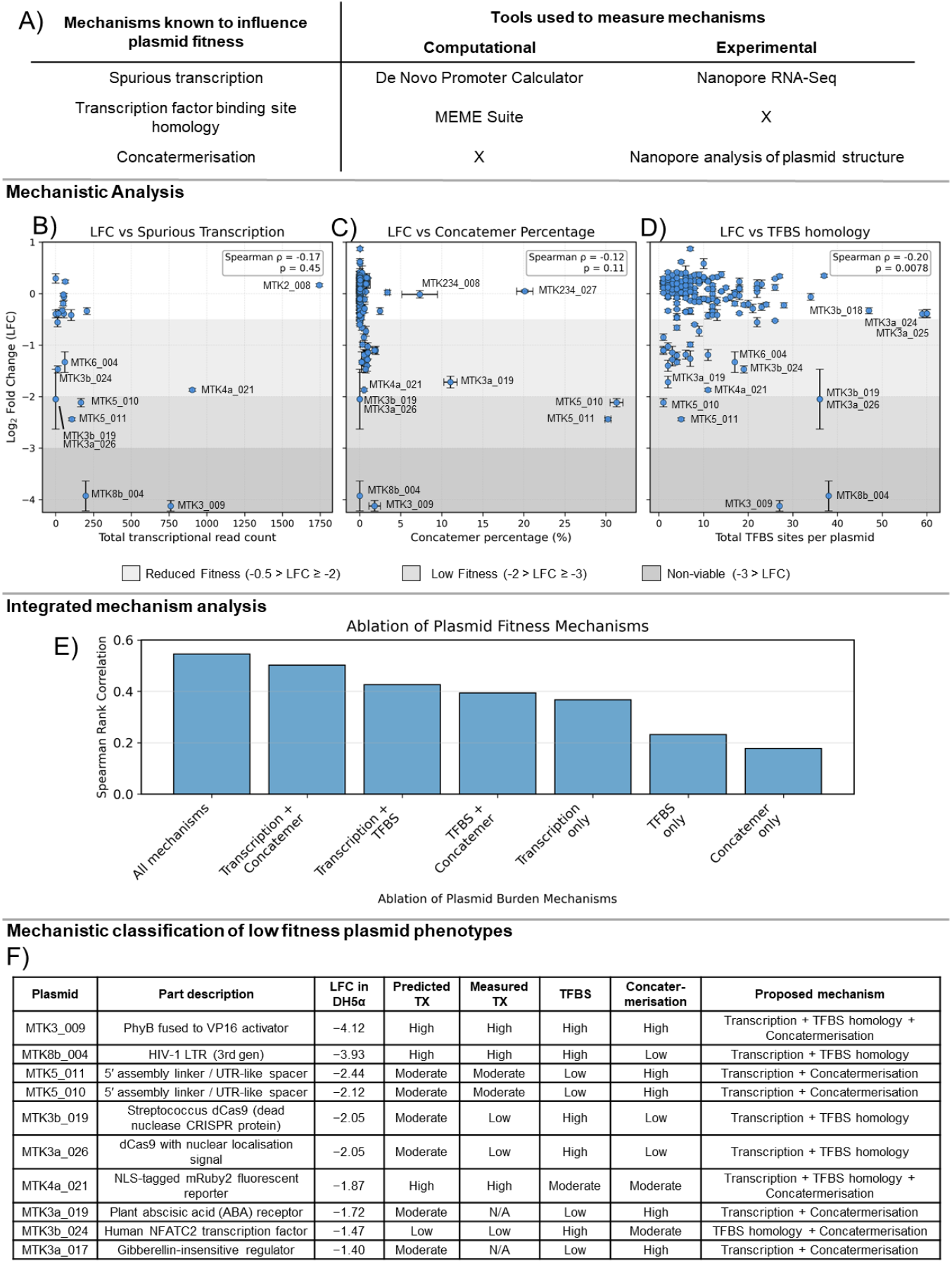
Multi-modal characterisation of plasmid sequence features relevant to fitness. A) Summary of plasmid–host interaction mechanisms analysed and corresponding measurements: predicted forward transcription, concatemerisation from Nanopore profiling, transcription-factor binding sites (TFBS) from motif analysis, and plasmid fitness (log fold change, LFC) from pooled growth assays. B) Relationship between targeted long-read transcription and plasmid fitness in *E. coli* DH5α. Total transcription represents the number of full-length Nanopore cDNA reads per plasmid, including forward and reverse transcripts. Each point represents a plasmid; vertical error bars indicate the standard deviation of LFC across three biological replicates. C) Plasmid fitness versus concatemer percentage for the same plasmids. Each point represents a plasmid; error bars show the standard deviation of LFC and concatemer percentage across three biological replicates. D) Plasmid fitness versus TFBS counts. Each point represents a plasmid; vertical error bars indicate the standard deviation of LFC across three biological replicates. E) Pairwise ranking ablation analysis showing the contribution of mechanistic feature groups to plasmid fitness prediction. Bars show Spearman correlation between measured and predicted LFC using different feature sets. F) Table of MTK plasmids, including function, LFC in DH5α, and inferred mechanism of interaction.

Previous studies have shown that eukaryotic and viral DNA can contain sequences resembling σ^70^ −10/−35 motifs that are recognised by *E. coli* and induce spurious transcription.^(12-16)^ Therefore, we first assessed whether promoter-like sequences could explain the low plasmid LFC in the library. We estimated the total σ^70^-like transcriptional potential across the MTK library using the De Novo Promoter Calculator,^(32)^ which showed a modest negative correlation with plasmid fitness (r = −0.17, p = 0.02; **Fig. S4**). From this we selected twenty-four plasmids (**Table S1**) spanning the full predicted transcriptional strength versus plasmid fitness space for analysis using RNA sequencing (RNA-seq). To evaluate whether transcriptional activity itself was associated with reduced propagation, we quantified total RNA output for each of the selected plasmids and compared it to their fitness. Although the total transcriptional output did not correlate with plasmid fitness (ρ = −0.17, p = 0.45; **Fig. 2B**), the lowest fitness plasmids were disproportionately associated with high transcriptional output. Plasmids with moderate or high fitness, by contrast, showed wide variability in transcription levels. Therefore, elevated transcription is enriched among the plasmids with the lowest fitness but is not by itself a universal determinant of plasmid fitness.

To understand why some highly transcribed plasmids propagated poorly while others remained unaffected, we examined strand-resolved transcriptional profiles for each plasmid (**Fig. S5**). These coverage maps revealed that the position and orientation of transcription initiation, rather than total transcript abundance, determined whether transcription imposed a fitness cost. For example, MTK8b_004, which contains a strong HIV-1 LTR promoter, and MTK3_009, encoding the potent PHYB–VP16 transcriptional activator with a nuclear localisation sequence, showed extensive forward transcription across the plasmid backbone and consistently low fitness, consistent with transcription–replication interference. By contrast, MTK6_004, MTK4a_018 and MTK3b_024 carried σ^70^-like motifs within their mammalian inserts but produced little to no detectable transcription and maintained near-neutral fitness. Finally, MTK2_007 (human pEF1α promoter) and MTK2_008 (human pUbc promoter) generated substantial RNA in *E. coli* despite being eukaryotic promoters, yet both plasmids propagated efficiently. These observations suggest that recognition of mammalian regulatory elements by σ^70^ is not inherently detrimental and that elevated transcription is sufficient to explain poor fitness in some cases but is not a reliable predictor across the library. Differences in where and how transcription is initiated therefore emerge as candidate factors underlying divergent outcomes.

These results resolved plasmids whose low fitness arose from transcriptional interference but also revealed constructs with severe fitness defects in the absence of detectable transcription. Using the long-read sequencing data generated from the original pooled parallel assay, we examined plasmid topology and found clear evidence of structural instability. Several low-fitness plasmids, including MTK3a_019, which contains a repeated receptor domain module, and MTK5_010 and MTK5_011, which each contain paired insulator elements arranged in direct orientation, accumulated extensive head-to-tail concatemers (**Fig. 2C**). These multimerised forms are known to interfere with replication fork progression and impair segregation, providing a structural explanation for reduced maintainability independent of transcriptional output.(^21, 22^)

A subset of low-fitness plasmids remained unexplained by either transcription or structural instability. Prior work has shown that plasmids can impose burden by sequestering global transcription factors, thereby altering chromosomal gene expression.(^19, 20^) Because mammalian regulatory elements often contain short motifs similar to *E. coli* regulator binding sites, we used MEME Suite to map homologous sites for 68 *E. coli* transcription factors across the MTK library.(^33, 34^) These binding sites spanned major functional transcription factor (TF) classes, with global regulators and stress factors showing the strongest shift toward reduced fitness, whereas metabolic, two-component, and pathway-specific TFs exhibited far weaker associations (**Fig. S6**). When comparing the ten lowest- and ten highest-fitness plasmids, low-fitness constructs generally exhibited higher TFBS load (**Fig. 2D**), consistent with regulatory factor sequestration. This included MTK3_009 and MTK8b_004, which displayed both high TFBS density and strong fitness defects. Crucially, MTK5_010 and MTK5_011 remained low fitness despite low TFBS counts, reinforcing that their burden arises from concatemerisation rather than regulatory sequestration. Our analysis thus suggests that TF binding site load contributes to plasmid burden in a defined subset of constructs, complementing rather than replacing transcriptional and structural mechanisms.

To place these mechanisms in a unified framework, we next asked how strongly each contributes to fitness when considered together. Because plasmid fitness in our pooled assay is defined relatively, plasmids increase or decrease in abundance compared to one another, we developed a pairwise ranking model to infer which of two plasmids is more likely to propagate successfully in *E. coli* (**Fig. S7**). This approach captures the logic of pooled competition directly, as rather than predicting absolute fitness values, the model learns the features that drive relative fitness advantage. Feature ablation demonstrated that the full model, incorporating transcriptional output, concatemer load and TFBS content, achieved the highest predictive performance (**Fig. 2E**). Removing any single feature class reduced performance, and models relying on only one mechanism performed substantially worse. Notably, models containing any two mechanisms recovered most of the predictive power of the full model, indicating that plasmid behaviour reflects interdependencies between transcriptional load, structural stability and regulator availability.

These results show that mammalian regulatory elements can interact with the *E. coli* host through multiple, partially overlapping mechanistic routes, spurious transcription, concatemerisation and transcription factor sequestration, and that these mechanisms act in combination to determine whether plasmids propagate efficiently, impose burden or fail to maintain themselves (**Fig. 2F**). This establishes the foundation of a plasmid–host interaction map linking specific sequence features to physiological outcomes and provides a framework for predictive, host-aware plasmid design.

### Contextual Effects in Multi-Part Mammalian Expression Cassettes

The single-part screen demonstrated that mammalian regulatory elements can impose transcriptional, structural, or regulatory burden in *E. coli*, but plasmids used in practice do not consist of parts in isolation. Instead, these elements are assembled into multi-part expression cassettes, where promoters, coding sequences, tags, and terminators sit in close proximity. In such architectures, transcription start sites can shift, secondary structures can interact, and protein-binding motifs can overlap. Therefore, the mechanistic effects observed for isolated parts may not be preserved when parts are combined. They may be buffered, amplified, or reconfigured entirely. The key question was thus whether the liabilities identified in the single-part screen persist in multi-part constructs, and whether combinations of parts produce new burden phenotypes that cannot be inferred from the parts alone.

To test this, we assembled a pooled library of four-part mammalian expression cassettes using the MTK. The library consisted of 8 promoters, 8 CDS, 8 localisation or regulatory tags, and 4 polyA elements, yielding 2,048 theoretical combinations, of which 1,847 (90.2%) were successfully recovered (**Fig. 3A**). This library provided a systematic framework to examine how the transcriptional, structural, and regulatory mechanisms identified in single parts manifest, or change, when parts are combined into operational cassettes. Following the same validated workflow established above, plasmid fitness was quantified as LFC in relative plasmid abundance over a 16-hour batch culture in DH5α. Fitness values spanned the full dynamic range of the assay (**Fig. 3B**).

**Figure 3.**
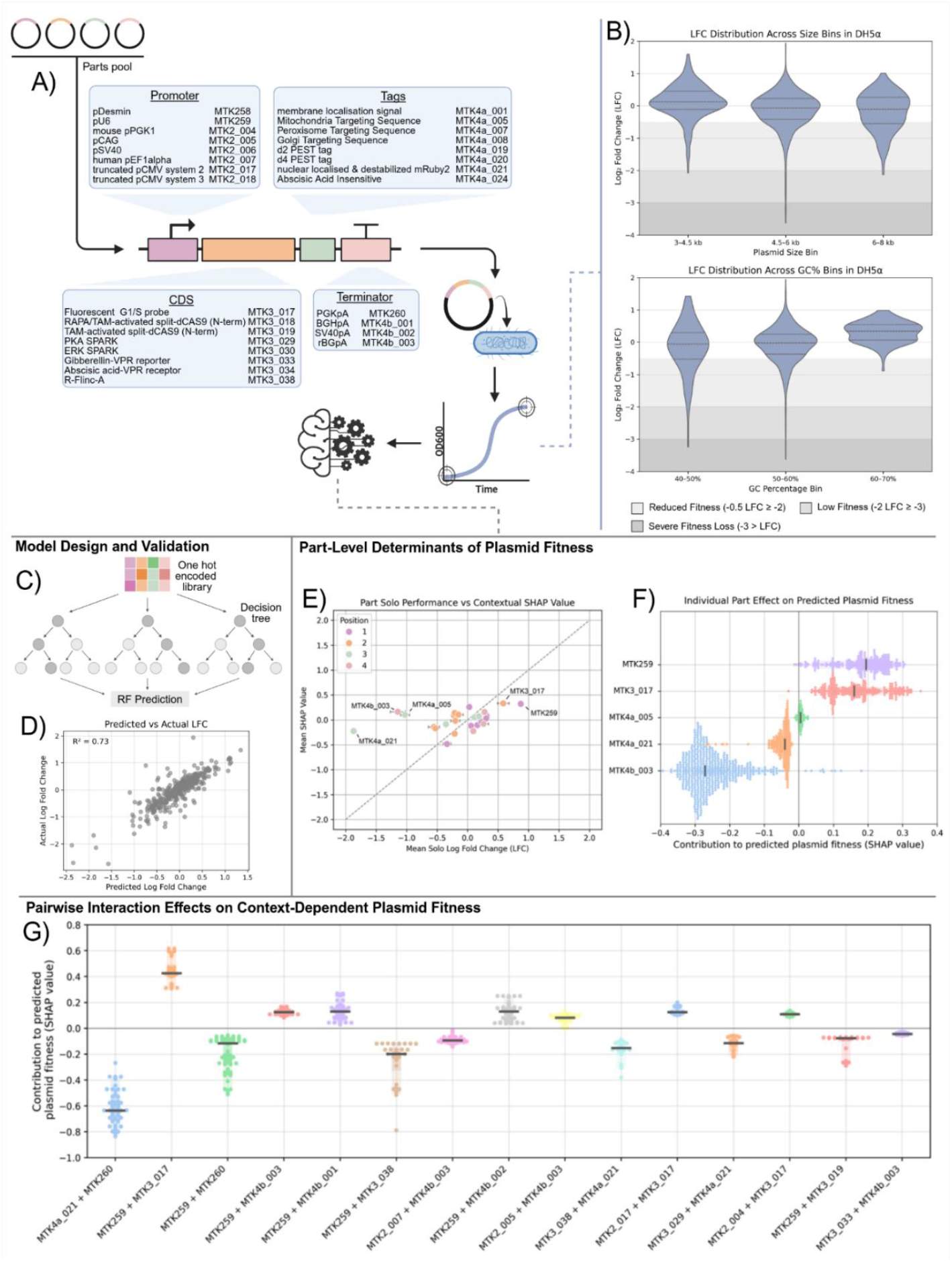
Combinatorial assembly and contextual performance of Mammalian Tool Kit (MTK) parts in *E. coli*. A) Four-part expression cassette assembly from the MTK library, comprising a promoter, coding sequence (CDS), localisation tag, and terminator, combined by one-pot Golden Gate cloning and propagated in *E. coli* DH5α. Pooled fitness was measured by quantifying plasmid abundance over a 16-hour batch culture. B) Plasmid log fold change (LFC) plotted against plasmid size (top) and GC content (bottom) for 1,847 assembled constructs. Each point represents one plasmid. C) Overview of the random forest regression workflow used to predict plasmid fitness from one-hot-encoded part identities. D) Predicted versus measured plasmid LFC for the random forest model. Each point represents one plasmid. E) Comparison of mean part fitness (LFC) with contextual effect estimated by mean SHAP value for each part. Points are coloured by cassette position (promoter, CDS, tag, terminator). F) SHAP value distributions showing part contributions to predicted plasmid fitness across all assemblies. Each ridge represents one part. G) Pairwise SHAP interaction values between selected parts. Points represent individual assemblies; distributions summarise interaction variability across the library.

Consistent with the single-part screen, plasmid size and GC content showed statistically significant but modest associations with fitness in multi-part constructs (size: ρ = −0.27, p < 0.001; GC content: ρ = 0.25, p < 0.001; **Fig. S8**). However, stratifying plasmids by size or GC bins revealed substantial overlap in fitness distributions (**Fig. 3B**). Constructs in the 4.5–6 kb range displayed a noticeably heavier tail of severely low fitness plasmids than the 6–8 kb bin, indicating that size is not a strong factor in determining plasmid propagation. Plasmids with higher GC content were less likely to have reduced fitness, whereas assemblies in the 40– 60% GC range spanned the full dynamic range from neutral to severely impaired propagation. Although this pattern, does not establish that AT-rich assemblies intrinsically lead to reduced fitness, it is indicative of these sequences being more prone to lower plasmid fitness. An explanation for this, is their increased likelihood of forming motifs homologous to σ^70^ promoter elements and driving spurious transcription in *E. coli*.^(16-18)^

To determine whether plasmid fitness could be predicted from the identities of the assembled parts, we trained a random forest regression model in which each plasmid is represented solely by the identities of its four constituent parts (**Fig. 3C**). Random forests are appropriate in this setting because they are well suited for a dataset of this size and can capture non-linear and combinatorial relationships without assuming additivity. The model achieved strong predictive performance (R^2^ = 0.74; **Fig. 3D**), indicating that a substantial portion of plasmid propagation is encoded directly in part composition and arrangement, rather than arising from stochastic variation or batch-specific effects. Model residuals displayed no systematic bias (**Fig. S9**), indicating that the learned structure reflects real biological constraints.

To interpret how individual parts influence plasmid maintainability within assembled cassettes, we computed SHAP (SHapley Additive exPlanations) values, which quantify the contribution of each part to the model’s prediction for each construct. A positive SHAP value indicates that the part improves propagation, while a negative value indicates that it contributes to burden, always in the context of the parts it is paired with. Comparing each part’s single-part fitness to its mean SHAP value in multi-part assemblies revealed substantial context-dependence (**Fig. 3E**). For example, nuclear mRuby2 and a mitochondrial targeting tag, which imposed strong burden when present alone, were buffered in assemblies with promoters such as pCAG and terminators such as PGKpA, indicating that context can suppress transcriptional interference or reduce regulatory sequestration. Conversely, the pU6 promoter, which propagated efficiently as a single part, imposed strong burden in specific multi-part constructs, suggesting that otherwise benign promoters can become detrimental when transcriptional load is distributed across several adjacent regulatory junctions.

SHAP distributions across the full library revealed three categories of part behaviour (**Fig. 3F**). Consistent enhancers of maintainability included pCAG, pU6, and the G1/S cell-cycle reporter, which uniformly improved propagation across contexts. Consistently burden-inducing parts, such as EF1α and the rBGpA terminator, decreased plasmid fitness regardless of context, indicating intrinsically high interaction cost with the bacterial host. A third group, including the SV40 promoter, PKA-SPARK signalling reporter, and R-Flinc-A cAMP reporter, displayed strong context-sensitivity, generating burden only in specific promoter–terminator environments. These patterns mirror the mechanistic classes observed in single parts but demonstrate that context determines the extent to which those mechanisms are expressed.

Finally, to test whether parts interact non-additively, we quantified pairwise SHAP interaction values (**Fig. 3G**). Several strong interaction patterns emerged. PGKpA, neutral on average, became strongly detrimental when combined with PKA-SPARK or nuclear mRuby2, but beneficial when paired with the G1/S reporter. Similarly, pU6 switched from beneficial to detrimental depending on terminator selection. These results show that plasmid fitness in *E. coli* is an emergent property of part–part and part–host interactions, not the sum of individual part behaviours.

### A Sequence-Based Predictive Framework for Plasmid Manufacturing Performance

Our analyses revealed that plasmids containing mammalian and viral regulatory sequences can impose substantial cell burden in *E. coli*, leading to poor propagation efficiency and reduced yield. These foreign regulatory elements frequently contain cryptic σ^70^-like promoters, local repeat structures, and short motifs resembling binding sites for global *E. coli* transcription factors, any of which may interfere with the transcriptional, structural, or regulatory networks required for stable plasmid replication.(^16, 17, 19-21, 23^) In the multi-part expression cassette library, we showed that these effects were part- and context-dependent, and that plasmid burden could be modelled from part identity using a random forest approach. However, the model was constrained to the exact set of parts included in the training library and cannot evaluate newly introduced, synthetic, mutated, or externally sourced sequences, which increasingly form the basis of modern plasmid engineering. A genuinely generalisable design framework must therefore be able to infer manufacturing performance directly from primary DNA sequence, without relying on prior annotation or assumptions about part function.

To determine how primary DNA sequence alone influences plasmid propagation, we developed a variant of the pooled growth assay described above, adapted specifically to remove part identity and contextual effects. A library of plasmids carrying fully degenerate 192 bp inserts in a constant backbone was transformed and propagated in 100 mL batch culture in *E. coli* DH5α, using the same growth conditions as in the part screens. However, instead of whole-plasmid long-read sequencing, we used Illumina amplicon sequencing targeting the insert, as only relative insert abundance was required in this context and we acquired more reads. Plasmid abundance at the start and after 16 hours of growth was quantified, and LFC provided a direct measure of sequence-driven effects on propagation efficiency, independent of part identity, regulatory function, or plasmid architecture (**Fig. 4A**).

**Figure 4.**
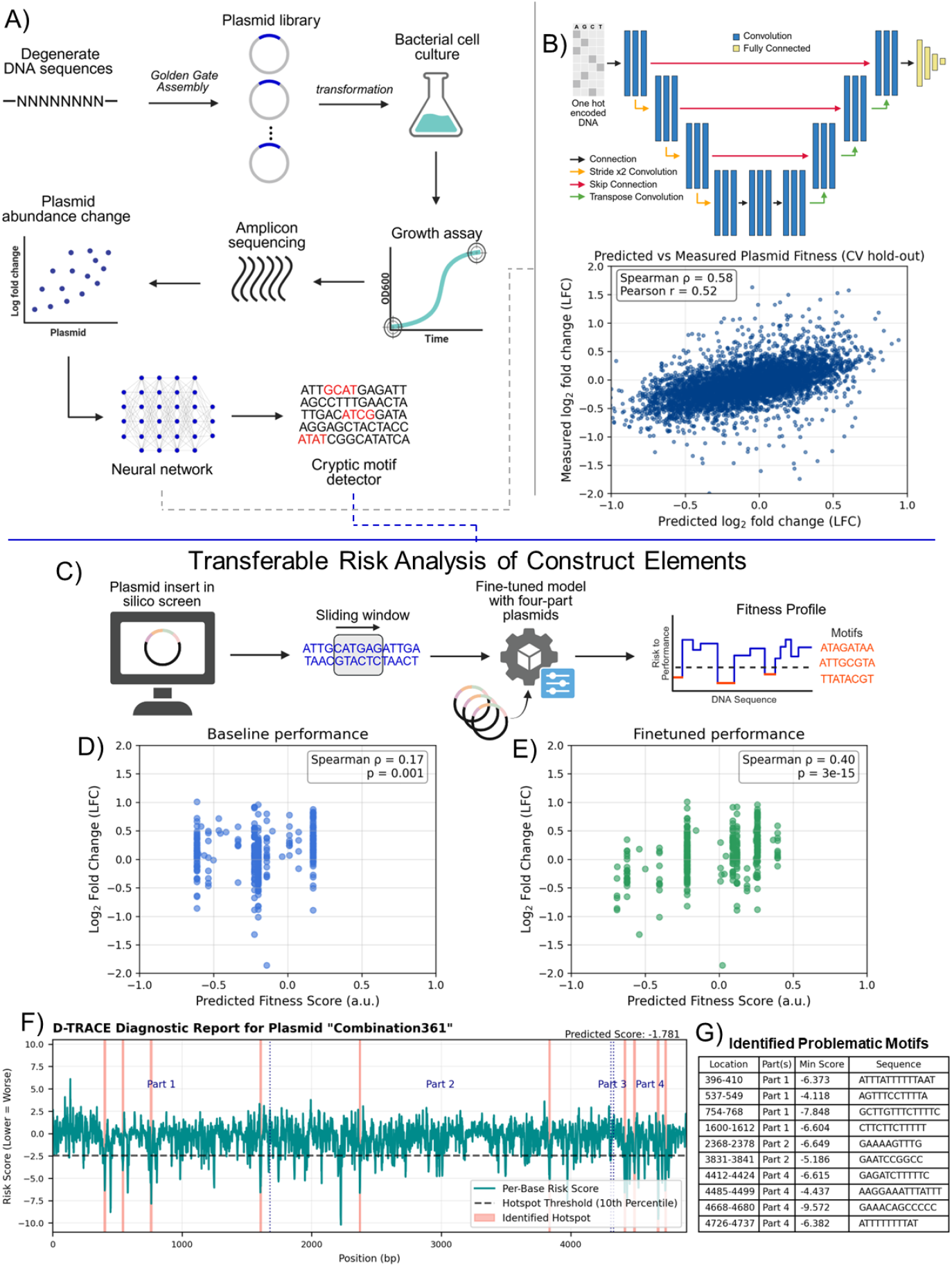
Deep learning framework for identifying sequence features influencing plasmid fitness. A) Workflow for the degenerate insert screen. A library of fully random 192 bp sequences was cloned into a constant plasmid backbone and propagated in *E. coli*. Plasmid fitness was quantified as log fold change (LFC) over batch culture, with abundance measured by amplicon sequencing. Fitness measurements were used to train a neural network on sequence features associated with plasmid propagation. B) 1D U-Net convolutional neural network architecture used to model the relationship between DNA sequence and plasmid fitness (top). Scatter plot comparing predicted versus measured LFC for held-out inserts (bottom). C) Sliding-window inference workflow for long constructs. Plasmid sequences were scanned in overlapping windows to generate a per-base fitness profile after model fine-tuning. D) Baseline model performance before fine-tuning on the four-part plasmid dataset. Scatter plot shows predicted versus measured LFC. E) Model performance after fine-tuning, showing improved correlation between predicted and measured fitness. F) Example TRACE diagnostic output showing per-base risk scores across a representative plasmid. Dashed lines indicate part boundaries; red regions exceed the risk threshold. G) Table listing high-risk sequence regions identified in (F), including coordinates, part identity, and underlying sequence.

To learn how local sequence features influence plasmid propagation, we developed a U-Net based convolutional neural network (CNN) predicts propagation efficiency directly from primary DNA sequence while preserving nucleotide-level spatial information (**Fig. 4B**). Its encoder–decoder architecture enables the model to detect short motifs (≈6–20 bp) associated with transcriptional interference, structural instability, or transcription-factor sequestration, while also integrating broader contextual patterns across the sequence. The training objective combined supervised prediction with reverse-complement consistency, enforcing the physical symmetry of DNA, and included a pairwise ranking term to better reflect the competitive dynamics of the pooled assay.

Training was performed on more than 23 million nucleotides, distributed across 124,339 experimentally measured variants. Despite containing no regulatory annotations and no contextual information beyond the 192 bp input window, the model learned a reproducible sequence grammar that generalised across the library. Under strict cross-validation, it achieved a Pearson correlation of r ≈ 0.52 and a Spearman correlation of ρ ≈ 0.57 between predicted and measured LFC (**Fig. 4B**). These metrics indicate that the network captures both the linear and rank-order components of the sequence–fitness relationship, demonstrating that a substantial proportion of the variance in plasmid propagation is encoded directly within local sequence features of the plasmid itself. The predictive signal from the plasmid sequence alone demonstrates that propagation efficiency reflects reproducible, sequence-encoded interactions between plasmid DNA and host machinery, rather than unpredictable physiological variation.

To understand what the model had learned, we applied Integrated Gradients to obtain per-base contribution scores across sequences. High-impact motifs were strongly enriched for binding signatures of globally acting transcription factors such as IHF and Lrp, which regulate nucleoid structure and central metabolism (**Fig. S10)**. Sequences predicted to strongly bind these regulators were systematically depleted during growth, consistent with transcription factor sequestration diverting regulators away from their chromosomal targets. Motifs predicted to engage narrower or condition-specific regulators had more modest or context-dependent effects. These observations directly reflect the mechanistic analyses above: the physiological cost of a sequence depends not simply on whether it contains a regulatory motif, but on which regulatory network that motif perturbs.

Having established that the model learns sequence-intrinsic features predictive of propagation efficiency, we integrated the trained U-Net into a design-stage predictive platform, TRACE (Transferable Risk Analysis of Construct Elements; **Fig. 4C**). TRACE accepts arbitrary DNA sequences and returns both a propagation performance score and a nucleotide-resolution burden profile, enabling sequence-level diagnosis of plasmid manufacturing risk before synthesis or cloning.

We first applied TRACE to full mammalian expression plasmids to evaluate whether the sequence grammar learned from short, context-free inserts generalises to complete constructs. Because the model operates on 192 bp inputs, we used a sliding-window approach across full-length plasmids, generating overlapping window predictions that were aggregated into a plasmid-level score. Despite never having been trained on multi-kilobase plasmids, TRACE achieved a Spearman correlation of ρ ≈ 0.17 across our library of 1,847 four-part constructs propagated in *E. coli* DH5α (**Fig. 4D**). Notably, this level of performance is substantially above baseline, given that the model was required to infer plasmid-level behaviour by integrating hundreds of local sequence evaluations, demonstrating that the learned sequence grammar is genuinely transferable across construct scale and architecture. To incorporate contextual dependencies, we fine-tuned TRACE on a subset of the four-part plasmid library. Fine-tuning improved predictive performance to ρ ≈ 0.40 on a strictly held-out test set enriched for previously unseen part combinations (**Fig. 4E**), demonstrating that local sequence motifs and higher-order plasmid context can be integrated within a unified predictive framework in DH5α.

TRACE also provides diagnostic interpretability. For each plasmid, the model outputs a nucleotide-resolution burden landscape, allowing identification of specific sequence segments likely to impose transcriptional interference, replication stress, or transcription factor sequestration. An example plasmid profile (**Fig. 4F**) illustrates how detrimental local motifs associated with poor plasmid fitness can be identified and curated in summary with the motif sequence and local fitness score (**Fig. 4G**). These results establish TRACE as a practical, mechanism-informed design tool for plasmid production in DH5α. Plasmid propagation efficiency is therefore not an unpredictable emergent property of host physiology, but a quantitative and interpretable sequence-encoded trait that can be predicted, diagnosed, and engineered to enable rational, host-aware plasmid design.

## Discussion

Plasmid DNA is foundational to modern biotechnology and therapeutic development, yet its propagation in *E. coli* is difficult to predict. Instability is typically treated as an unavoidable consequence of incorporating DNA elements that are foreign to *E. coli*, and when it arises it is handled in an ad hoc manner rather than through general design principles. Previous studies have noted that eukaryotic, viral or even highly repetitive sequences can interfere with *E. coli*’s replicational, transcriptional, and regulatory processes, but these reports have been fragmented and system-specific, and were not designed to assess plasmid propagation, leaving no general framework for understanding why certain plasmids propagate poorly while others remain stable. Here, using quantitative profiling across millions of nucleotides, we show that plasmid propagation is a sequence-encoded phenotype that can be measured at scale, mechanistically dissected, and predicted.

Using a pooled competition assay, we found that DNA elements foreign to *E. coli* vary widely in their ability to propagate, even when cloned into the same backbone and cultured under identical conditions. These differences cannot be explained by plasmid size, GC content, or canonical burden metrics alone, consistent with previous work demonstrating that global metabolic load is a poor predictor of plasmid maintainability.(^7, 9^) Instead, we found that propagation efficiency reflects sequence-encoded interactions between plasmid DNA and the bacterial transcriptional and replication systems, interactions that have been difficult to map using traditional low-throughput assays.

Across millions of nucleotides tested, three mechanistically distinct liabilities consistently account for low plasmid propagation. First, spurious transcription arises when AT-rich or viral/eukaryotic motifs mimic σ^70^ promoter elements, driving unintended transcription and imposing substantial transcriptional load on the host.^(13-17)^ Second, structural instability arises when repetitive or insulator-rich regions promote concatemerisation. Prior work has reported this behaviour in certain viral cassettes and high-copy backbones, but here we show it is widespread and sequence-specific across mammalian construct libraries.^(21-24)^ Third, regulatory interference results from plasmid-encoded motifs homologous to *E. coli* transcription factor binding sites, acting as decoys that sequester regulators such as CRP, FNR, or IHF and disrupting cellular processes.(^19, 20^) While each mechanism has been described individually, their combined contributions to plasmid propagation have not been resolved prior to this work. A key insight is that these liabilities are not interchangeable, as some plasmids are dominated by transcriptional interference, others by concatemerisation, and others by regulator sequestration. Mitigating one liability does not alleviate the others, explaining why no single strain modification, intervention aimed at reducing unintended transcription, or recombination-suppressed background has historically provided a universal solution for improving plasmid stability and propagation.(^13-18, 21, 23, 28, 35^)

Extending the assay to multi-part expression cassettes showed that plasmid fitness is highly context dependent. The effect of a part on propagation varied depending on the other parts present in the cassette, with some combinations reducing the detrimental effects seen when parts were tested in isolation and others revealing liabilities that were not apparent on their own. These results mirror observations in synthetic biology that show component behaviour is often non-additive, and here this principle extends to the cryptic sequence features embedded within mammalian and viral DNA.(^36, 37^) For example, a sequence containing a cryptic transcriptional start site might be less detrimental when positioned next to another part that carries an unannotated terminator-like feature, illustrating how contextual buffering can occur even when the underlying elements are not intentionally designed. Overall, propagation efficiency emerges from the combined properties of the cassette rather than from the intrinsic behaviour of individual parts.

This combinatorial behaviour reflects an underlying regulatory “grammar” unintentionally encoded across the plasmid sequence that can interact with *E. coli*. To capture and predict this grammar directly from DNA, we developed, a model that first learns general sequence rules from over a million nucleotides and then transfers this knowledge to structured plasmids via fine-tuning. TRACE predicts plasmid propagation performance directly from the sequence and highlights liabilities at base resolution. Unlike promoter-or motif-based predictors that rely on predefined regulatory features,(^32, 38, 39^) TRACE learns sequence–function relationships de novo, enabling it to capture liabilities that are cryptic, or only emerge in combinatorial contexts. Importantly, this predictive framework works across plasmid architectures and is not limited to a single backbone or part set.

These results establish a unified and mechanistically grounded framework for host-aware plasmid engineering. Rather than treating plasmid instability as an unavoidable artifact of biology, this work shows that sequence-encoded liabilities can be diagnosed systematically and predicted in silico. TRACE provides a practical route toward identifying cryptic promoter activation, transcription factor sequestration, and recombination-prone architectures before synthesis or cloning. While this study does not implement optimisation in closed loop, it establishes the analytical and predictive foundations required for rational design of manufacturable plasmids. In doing so, it converts plasmid propagation from an unpredictable bottleneck into a quantifiable engineering constraint, laying the groundwork for future design-build-test cycles in synthetic biology, mammalian expression, and therapeutic manufacturing.

## Methods

### Plasmid Library Construction

Single-part plasmid libraries were assembled using the Mammalian Tool Kit (MTK) modular cloning framework, sourced from Addgene.^(11)^ Individual regulatory elements (promoters, UTRs, coding sequences, localization tags, and polyadenylation sequences) were cloned into a common bacterial backbone containing a chloramphenicol resistance marker and a pMB1-derived origin of replication. Multi-part expression cassettes were assembled by one-pot Golden Gate Type IIS assembly using BsaI-HFv2 (NEB) and T4 DNA Ligase (NEB) according to the MTK assembly format, with reaction cleanup performed by SPRI bead purification. Fully degenerate 192 bp libraries were synthesized as pooled oligonucleotide libraries (Twist Bioscience) and cloned into the same backbone using Golden Gate assembly. All plasmid constructs were verified by Sanger sequencing or Nanopore plasmid re-basecalling prior to pooled assays.

### Bacterial Strains and Transformation

Chemically competent *E. coli* DH5α, DH10B, and Stbl3 (Thermo Fisher / NEB) were used for pooled plasmid propagation assays. Pooled plasmid libraries were transformed into each strain via electroporation, recovered for 1 h in SOC, and directly diluted into selective LB medium (chloramphenicol, 25 μg/mL). Cultures were propagated for 16 h at 37°C with orbital shaking (200 rpm). No colony bottlenecking steps were performed after transformation to maintain library diversity.

### Pooled Competitive Growth Assay and DNA Extraction

Plasmid pools were sampled at two time points: immediately after transformation recovery (t = 0) and after 16 h batch culture (t = 16 h). Plasmid DNA was extracted using a miniprep kit (Qiagen / Macherey-Nagel), following a modified protocol optimized for pooled plasmid yield. For copy-number determination and RNA extraction experiments, aliquots were taken directly from the same cultures used for sequencing-based fitness measurements.

### Long-Read Sequencing and Fitness Quantification

Plasmid abundance was quantified using Oxford Nanopore sequencing (MinION / Flongle flow cells). Libraries were prepared using the Ligation Sequencing Kit (SQK-LSK109) with native barcoding (EXP-NBD104). Reads were basecalled with Guppy (super-accuracy mode) and circularized plasmid sequences were identified using mappy (minimap2) alignment to the plasmid reference library. Relative abundance of each plasmid was computed as reads-per-construct normalized to total plasmid-mapping reads per sample.

Biological replicates were processed independently and combined by median LFC per construct.

### Plasmid Copy Number and Growth Measurements

Plasmid copy number per cell was quantified by qPCR using primer pairs targeting the replication origin and the chromosomal *dxs* locus as the single-copy reference gene. OD_600_ growth curves were acquired in 96-well format using a microplate reader (Tecan / BMG Labtech) to determine growth rate and maximum culture density. Plasmid yield was measured from purified plasmid preparations using BioDrop spectrophotometry.

### Concatemer Detection by Long-Read Sequencing

Structural plasmid forms were quantified by classifying long Nanopore reads according to the number of full-length plasmid units present. Continuous plasmid-repeat arrays (2x or more units per read) were counted as concatemers, and percentage concatemerisation per construct was computed relative to all reads aligning to that plasmid backbone.

### Targeted long-read transcriptomics

To monitor possible unwanted transcription generated from the mammalian part plasmids in E. coli cells, targeted long-read RNA-seq was used.

RNA was extracted from cells using the RNeasy Protect Bacteria Mini Kit (QIAGEN, 74524) following the manufacturer’s protocol. Briefly, collected cells were stabilised in RNAprotect Bacteria Reagent, pelleted, and stored at -70°C or processed immediately. Pellets were lysed using TE buffer containing 1 mg/mL lysozyme (Thermo Scientific, 90082) as per the Enzymatic Lysis of Bacteria protocol. The lysate was mixed with RLT buffer supplemented with β-mercaptoethanol (Sigma, 444203) and processed with RNeasy Mini Kit columns. On-column digestion was performed with the QIAGEN RNase-Free DNase Set (QIAGEN, 79254). RNA was eluted in 30 µL of UltraPure™ DNase/RNase-Free Distilled Water (Invitrogen, 10977035). RNA concentration was measured using a Biochrom BioDrop μLITE+ (Biochrom, 80300655), and integrity was checked by agarose gel electrophoresis.

Targeted cDNA synthesis was performed on up to 5 µg of RNA using the Template Switching RT Enzyme Mix (NEB, M0466). Custom reverse transcription primers TC_PR_109 (for reverse-direction transcripts) and TC_PR_110 (for insert-direction transcripts) were used for targeted synthesis, along with the template switching oligo TC_PR_113. Second-strand synthesis was carried out using Q5 High-Fidelity DNA Polymerase (NEB, M0491) with dNTPs, Q5 Reaction Buffer, and RNase H (NEB, M0297). The resulting cDNA was purified using AMPure XP Reagent (Beckman, A63881) and eluted in 11 µL UltraPure™ Water.

Sequencing libraries were prepared from the cDNA pools using the Native Barcoding Kit 24 V14 (SQK-NBD114.24, ONT) following the standard supplier protocol. These libraries were loaded onto R10.4.1 flow cells (FLO-MIN114, ONT) and sequenced using a MinION Mk1D device for 72 hours. The raw POD5 data produced was basecalled and de-multiplexed using Dorado (v1.0.2) and the ‘dna_r10.4.1_e8.2_400bps_sup@v5.2.0’ model.

To ensure that only full-length targeted transcripts were included, nanofilter (v1.0) was used to extract reads incorporating a targeting primer sequence on one end (sequences from TC_PR_109 for reverse-direction transcripts or TC_PR_110 for insert-direction transcripts) and the strand switching oligo sequence (from TC_PR_113) on the other. After filtering, reads were mapped using minimap2 (v2.30-r1287) to the full set of possible plasmid sequences, with no secondary alignments. Read depth profiles across each plasmid were then generated using the samtools (v1.22.1) ‘depth’ command and the resultant profiles visualized using custom Python (v3.11) scripts with numpy (v2.3.4), matplotlib (v3.10.7), and biopython (v1.86).

### Motif and Promoter Analysis

Promoter-like sequence activity was predicted using a bacterial sigma70 promoter scoring model. TFBS-like motifs were identified using MEME Suite and matched to *E. coli* TF-binding motifs using Tomtom against DPInteract motif database. Motif significance was assessed using E-value < 1e–4 and FDR-corrected q-values where applicable.(^33, 40^)

### Statistical Analysis

Correlations were computed using Spearman rank (ρ) or Pearson (r) coefficients. Multi-strain comparisons were performed by paired or unpaired rank tests as appropriate. All statistical tests were two-sided unless stated otherwise. No samples, data, or replicates were excluded.

### Machine Learning and Feature Attribution

DNA sequences were one-hot encoded. Propagation efficiency for the 192 bp library was predicted using a U-Net–based 1D convolutional neural network with an encoder–decoder architecture that preserves nucleotide-level spatial information. The model incorporated reverse-complement consistency and a pairwise ranking term in addition to mean-squared error to reflect the competitive nature of the pooled assay. Training was performed on 124,339 experimentally measured variants. Model accuracy was assessed by Pearson and Spearman correlation on held-out cross-validation folds.

For multi-kilobase plasmids, predictions were obtained using sliding-window inference (192 bp window, stride defined in text). Outputs from five independently trained U-Net models were averaged to obtain consensus predictions. Base-resolution contribution scores were computed using Integrated Gradients applied to the ensemble. High-scoring regions were extracted and provided to MEME for de novo motif discovery. Identified motifs were matched to *E. coli* transcription factor binding motifs using Tomtom against the DPInteract database (E-value < 1e-4).

## Acknowledgements

The authors would like to thank Michael Crone (Full Circle Labs, London) for support and advice on the nanopore sequencing carried out in this work. This work was supported by the EPSRC Centre for Doctoral Training in BioDesign Engineering (EP/S022856/1) (to T.C., J.M. E.T. and F.C.). F.C. was also partly funded by the Bezos Earth Fund through the Bezos Centre for Sustainable Protein (BCSP/IC/001), the UK National Alternative Proteins Innovation Centre (NAPIC), which is an Innovation and Knowledge Centre funded by the Biotechnology and Biological Sciences Research Council (BBSRC) and Innovate UK (BB/Z516119/1). F.C., T.E.G. and J.H.C.F. were partly funded by the Engineering and Physical Sciences Research Council under the EEBio Programme Grant (EP/Y014073/1). T.E.G. was supported by Royal Society University Research Fellowship (URF\R\221008).

## Author contributions

T.C, T.E. and F.C. designed the project. T.C. performed the main experiments and computational analysis. J.H.C.F. performed the RNA extractions and cDNA synthesis for the random transcripts analysis. J.M. advised on the computational framework. T.E.G. performed nanopore sequencing and computational analysis for the random transcript experiments. T.C, F.C. and T.E. wrote the manuscript. All authors read and edited the manuscript.

## Data Availability Statement

All source data and plasmid maps for the data sets presented in the manuscript are available with this publication as related manuscript files. Constructs are available upon request.

## Software availability statement

The modelling code developed in this study is available at https://github.com/tcopeman21/plasmid-host-interactions.git

