## Supplementary Information for "Quantitative profiling of millions of nucleotides reveals sequence-encoded interactions that govern plasmid propagation"

### Supplementary Figures

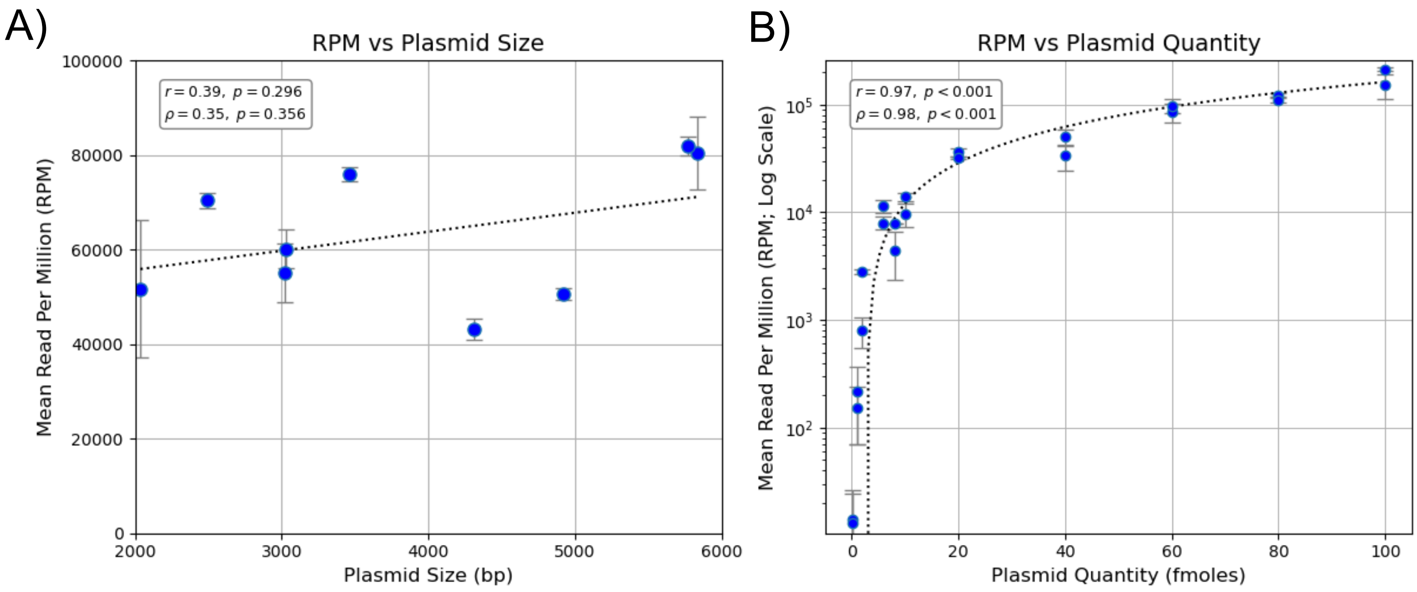


**Figure S1. Validation of Oxford Nanopore sequencing for relative plasmid abundance quantification.**

A) Normalised read counts (reads per million; RPM) were quantified from an equimolar pool of plasmids spanning 2–6 kb to assess size-dependent bias in sequencing. RPM showed a weak, non-significant association with plasmid size (Pearson $r = 0.39$, $p = 0.296$; Spearman $\rho = 0.35$, $p = 0.356$). B) Normalised read counts (RPM) were quantified from a pool of similarly sized plasmids prepared at increasing femtomole quantities to assess quantitative linearity. RPM scaled strongly with input plasmid quantity across more than two orders of magnitude (Pearson $r = 0.97$, $p < 0.001$; Spearman $\rho = 0.98$, $p < 0.001$). Error bars represent the standard deviation across independent replicates ($n$ = 3).

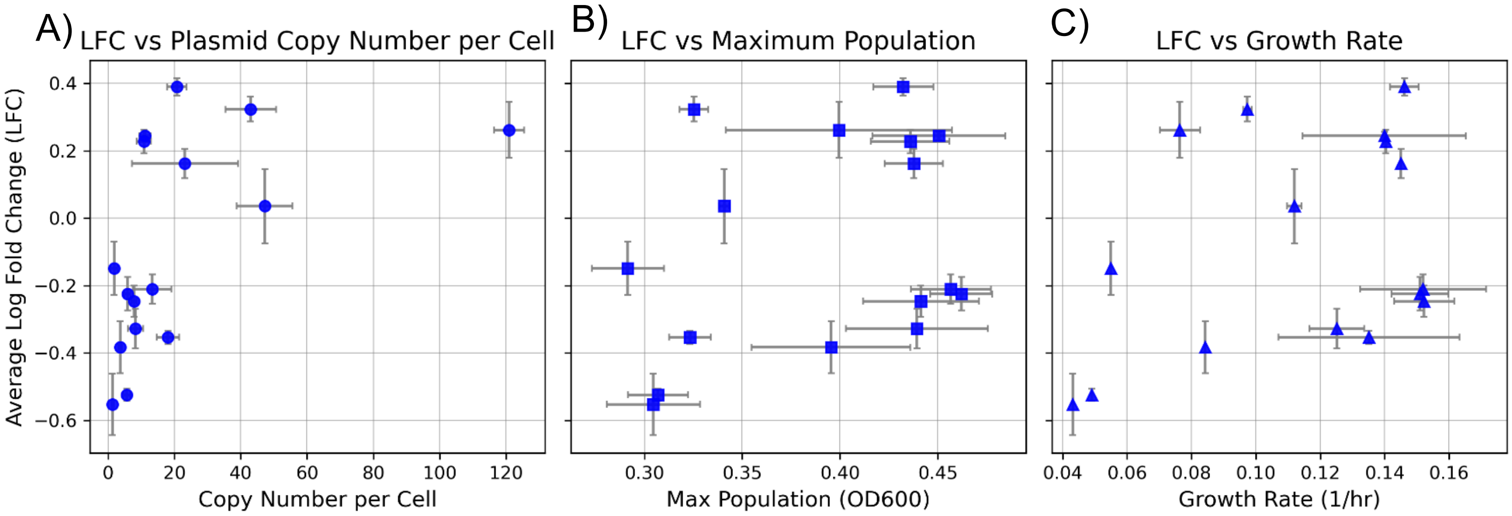


**Figure S2. Correlation of pooled parallel assay LFC with standard plasmid and host metrics.**

The panels correlate the average log fold change (LFC) from the pooled parallel assay with: (A) plasmid copy number per cell (Spearman r = 0.71, p = 0.002), (B) maximum population density (OD600) (r = 0.24, p = 0.368), (C) *E. coli* growth rate (r = 0.29, p = 0.274). Error bars represent the standard deviation across biological replicates (*n* = 3).

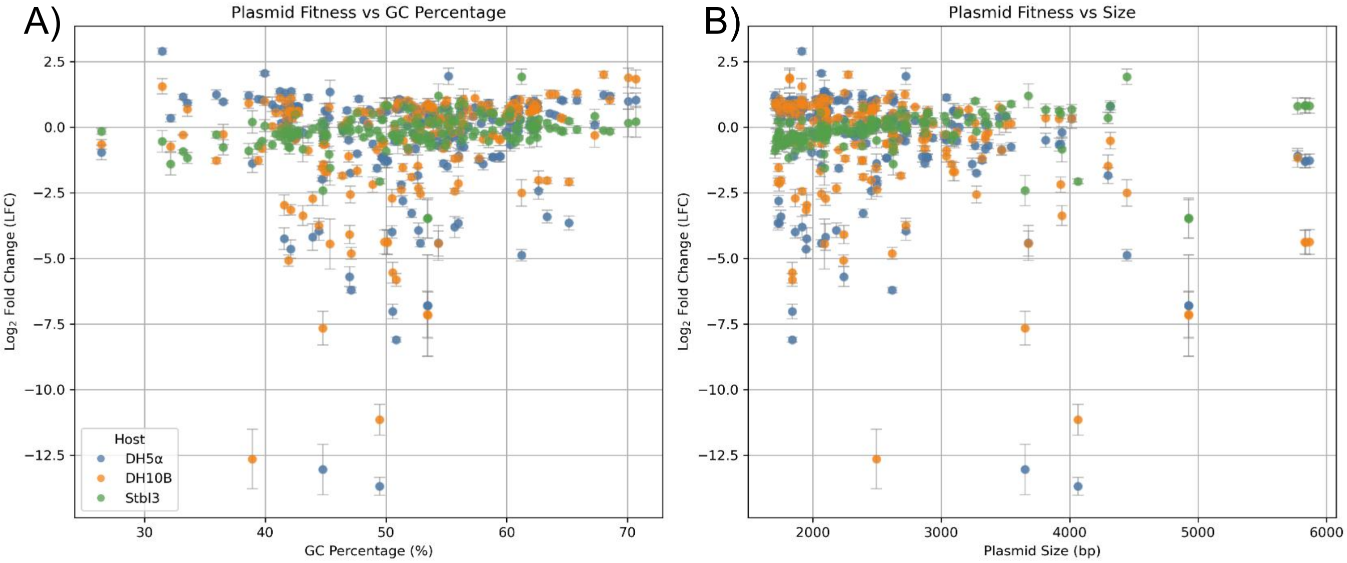


**Figure S3. Relationship between plasmid fitness, GC content, and plasmid size across *E. coli* hosts.**

Plasmid fitness, quantified as $\log_2$ fold change (LFC) is shown as a function of (left) GC percentage and (right) plasmid size. Each point represents an individual plasmid measured in three *E. coli* hosts: DH5α (blue), DH10B (orange), and Stbl3 (green). Error bars indicate the standard deviation across independent biological replicates ($n$ = 3).

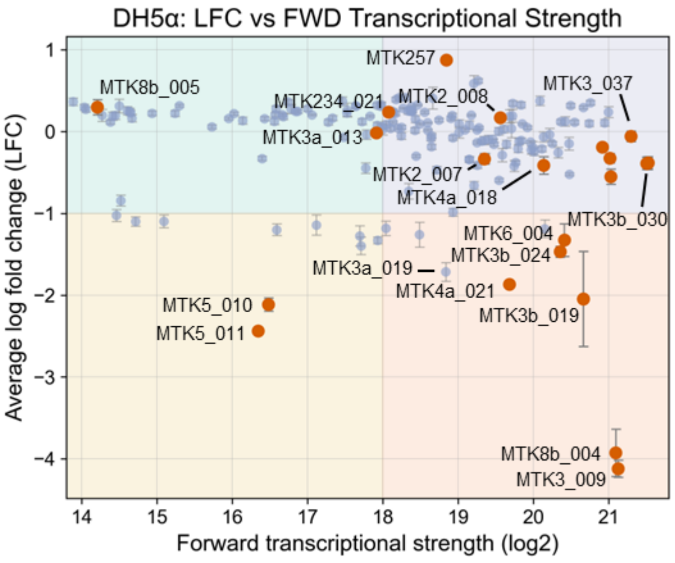


**Figure S4. Relationship between predicted σ⁷⁰-like forward transcriptional strength and plasmid fitness in DH5α.**

Scatter plot comparing forward transcriptional strength ($\log_2$ predicted $\sigma^{70}$-like promoter activity) to plasmid fitness (average log fold change, LFC) across the MTK plasmid library in *E. coli* DH5α. Blue points represent all library plasmids (n = 192); orange points denote the twenty plasmids selected for RNA-Seq analysis. Although predicted transcriptional strength and fitness show no global correlation (Pearson r = −0.171), the lowest-fitness constructs are disproportionately associated with high predicted transcriptional activity. Several plasmids with extensive predicted forward transcription, including MTK8b_004 (HIV-1 LTR promoter) and MTK3_009 (PHYB–VP16 activator), show marked fitness defects, whereas mammalian promoters MTK2_007 (pEF1α) and MTK2_008 (pUbc) exhibit substantial predicted transcription yet maintain near-neutral fitness. Error bars indicate standard error of the mean (SEM) across biological replicates. These patterns highlight that promoter-like sequence content is enriched among low-fitness plasmids but is not itself sufficient to determine plasmid maintainability. All data are N=3.

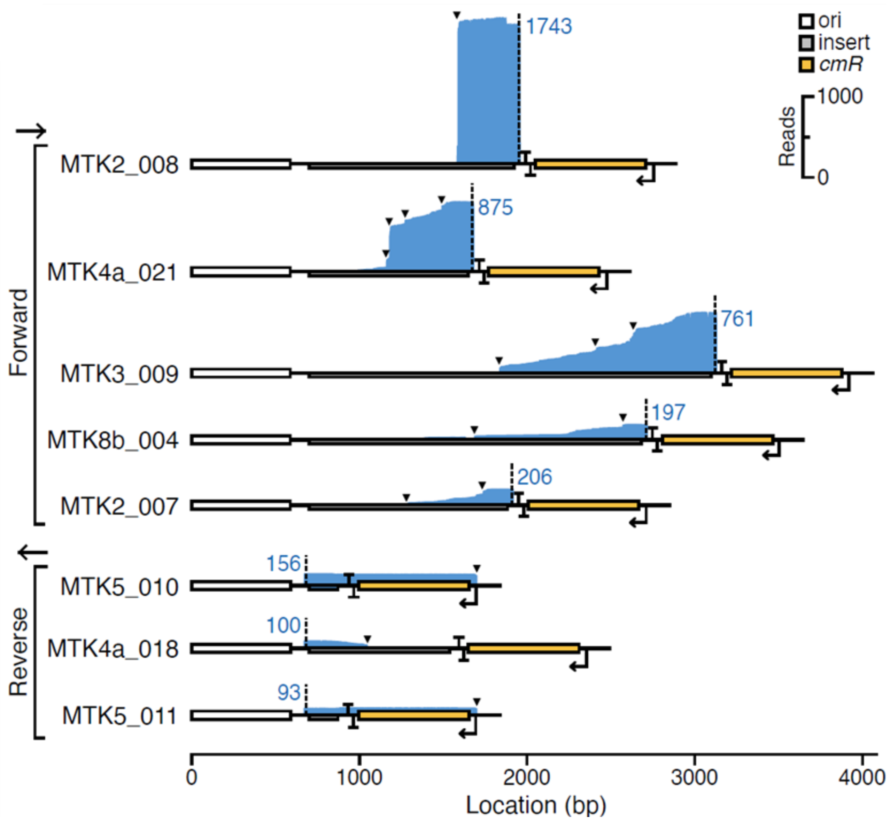


**Figure S5. Strand-resolved Nanopore RNA-Seq profiles reveal context-dependent transcription across selected plasmids.**

Forward and reverse strand RNA-Seq coverage across seven representative plasmids, illustrating distinct transcriptional behaviours underlying their fitness outcomes. Coverage (blue shading) is plotted across plasmid maps annotated with the replication origin (*ori*), mammalian insert, and *cmR* cassette. Plasmids MTK8b_004 and MTK3_009 exhibit extensive forward transcription traversing backbone regions, consistent with transcription–replication interference and their strongly reduced fitness. MTK4a_021 and MTK4a_018 display moderate but spatially restricted transcription, while MTK2_007 (pEF1α promoter) and MTK2_008 (pUbc promoter) show substantial RNA production despite maintaining near-neutral fitness, indicating that cross-recognition of mammalian promoters by $\sigma^{70}$ is tolerated when initiation occurs in non-disruptive regions. In contrast, low-fitness constructs MTK5_010 and MTK5_011 show predominantly reverse-strand transcription at low levels, reflecting that their fitness defects arise from structural instability rather than transcriptional load. Coverage values (dashed labels) indicate peak read counts per plasmid. These strand-resolved profiles highlight that transcriptional architecture, rather than total transcript abundance, determines whether spurious transcription imposes burden.

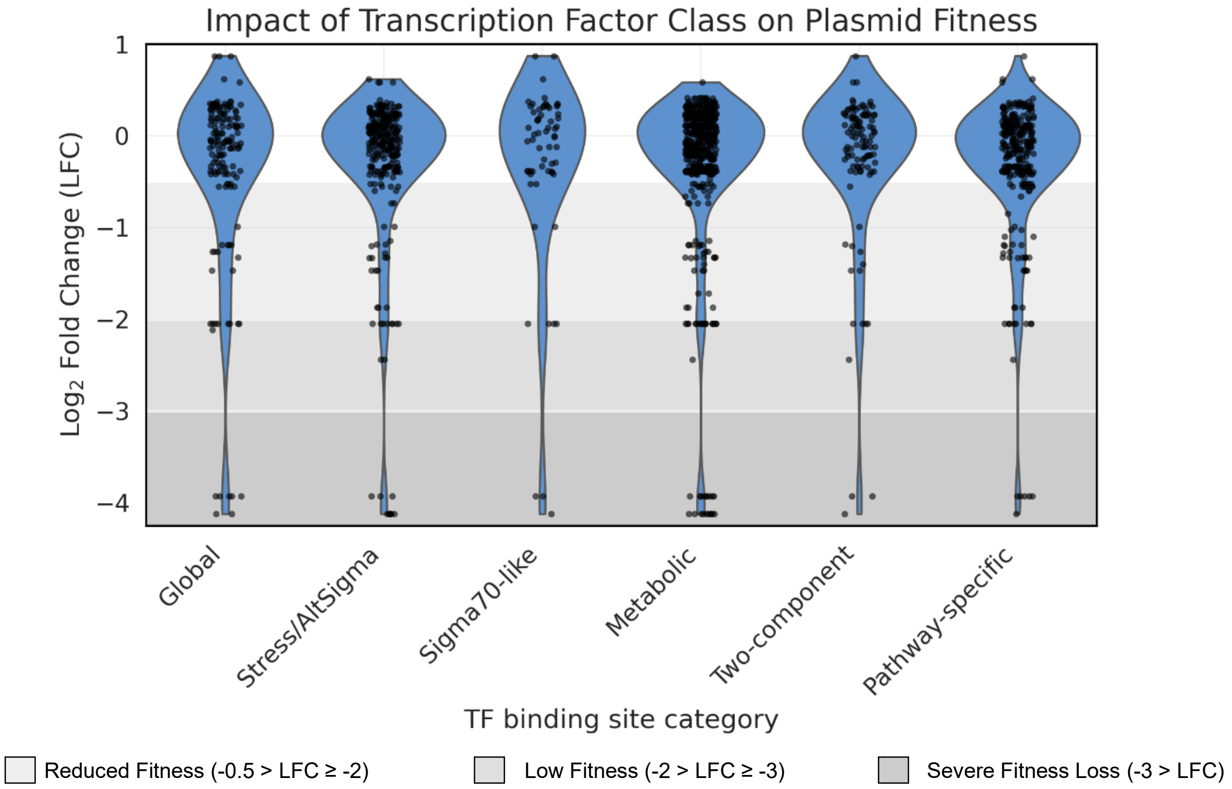


**Figure S6. transcription factor class on plasmid fitness.**

Violin plots of the $\log_2$ fold change (LFC) distribution for plasmids carrying at least one binding site for transcription factors (TFs) in each functional regulatory class. TF binding sites were identified from FIMO scans and assigned to six mechanistic categories: global regulators and nucleoid-associated proteins ("Global"), stress-response and alternative sigma factors ("Stress/AltSigma"), $\sigma^{70}$-like promoter fragments ("Sigma70-like"), metabolic regulators involved in carbon and amino-acid metabolism ("Metabolic"), non-global two-component response regulators ("Two-component"), and all remaining regulators with more pathway-specific roles ("Pathway-specific"). Each data point represents a plasmid–TF pair with ≥1 binding site in that class, so plasmids with sites for multiple TF classes can contribute points to multiple violins. Grey horizontal bands indicate the plasmid fitness ranges, including reduced fitness (light grey; -0.5 > LFC ≥ -2), low fitness (grey; -2 > LFC ≥ -3), and severe fitness loss (dark grey; -3 > LFC).

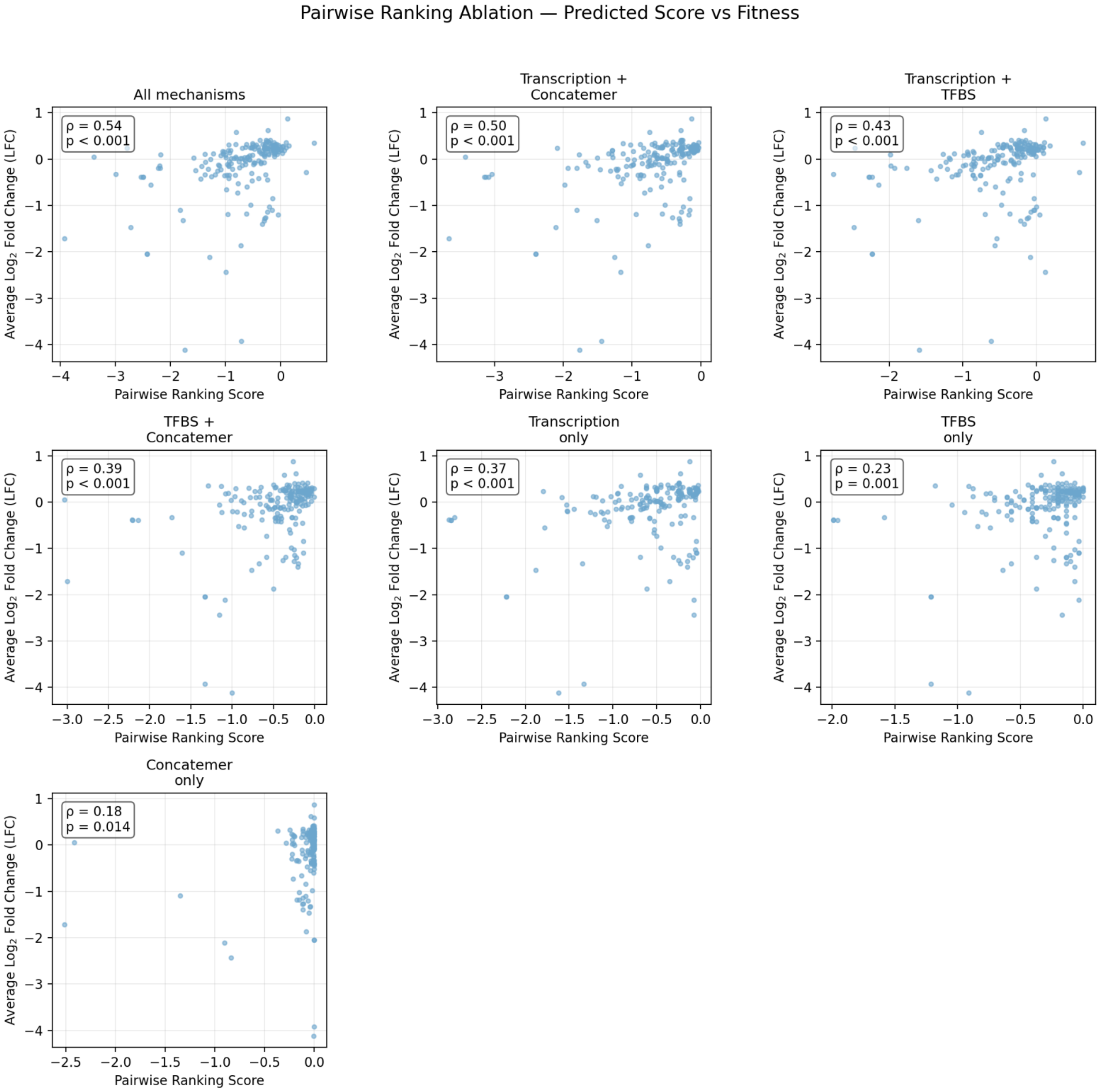


**Figure S7. Pairwise ranking ablation analysis of plasmid fitness mechanisms.**

The relationship between predicted pairwise ranking scores and plasmid fitness, measured as average $\log_2$ fold change (LFC), for models trained on different mechanistic feature information. Each panel represents an ablation in which predicted transcriptional burden (TX), predicted total transcription factor binding sites (TFBS), or concatermerisation frequency were included individually or in combination. Predictions were generated using a logistic pairwise ranking model trained on 60,000 random plasmid pairs per ablation. Spearman correlation coefficients (ρ) and associated p-values are reported in each panel. Models incorporating multiple mechanistic features achieved the strongest correspondence, with all mechanisms achieving ρ = 0.54, while single-feature models showed lower ρ correlation, consistent with plasmid propagation being influenced by multiple interacting sequence-encoded mechanisms.

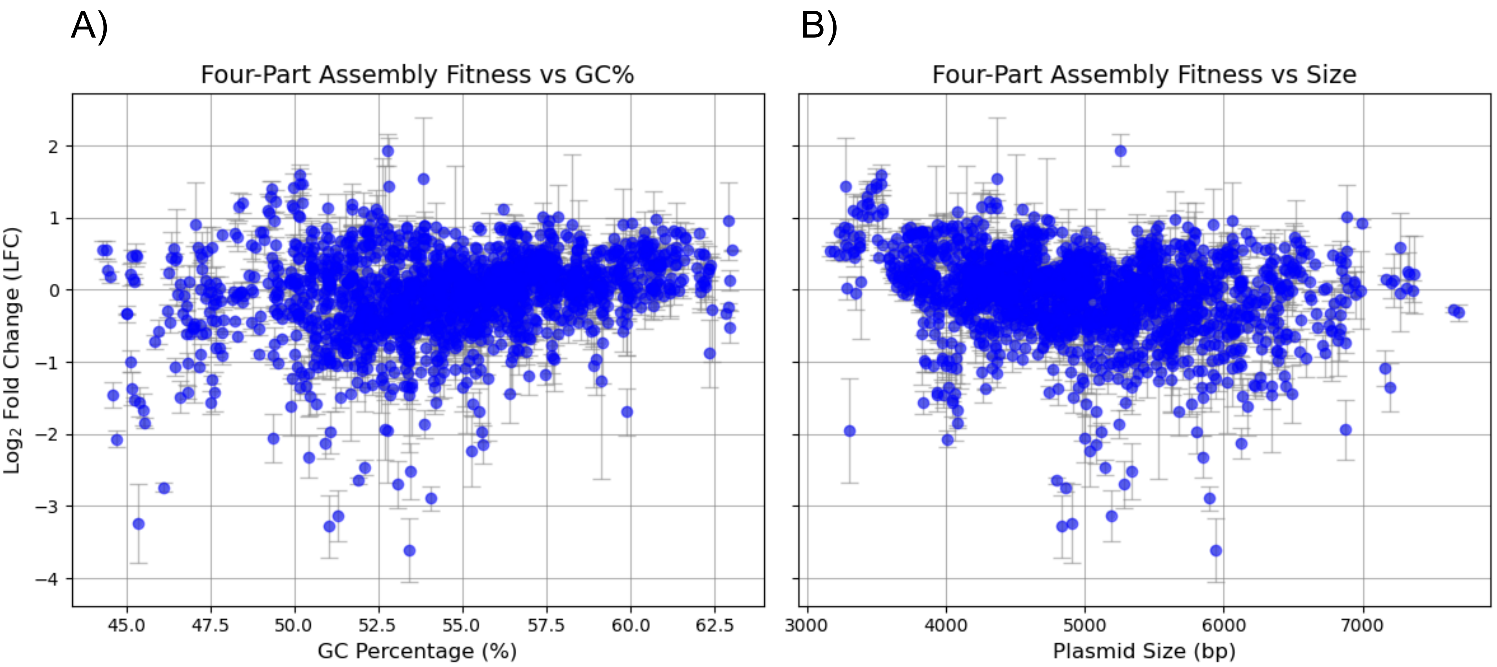


**Figure S8. Comparison between plasmid log fold change (LFC), plasmid size and GC content.**

A) Plasmid LFC GC percentage for four-part mammalian expression constructs propagated in *E. coli* DH5α. B) plasmid fitness versus total plasmid size. Each point represents an individual plasmid; error bars indicate standard deviation across three biological replicates.

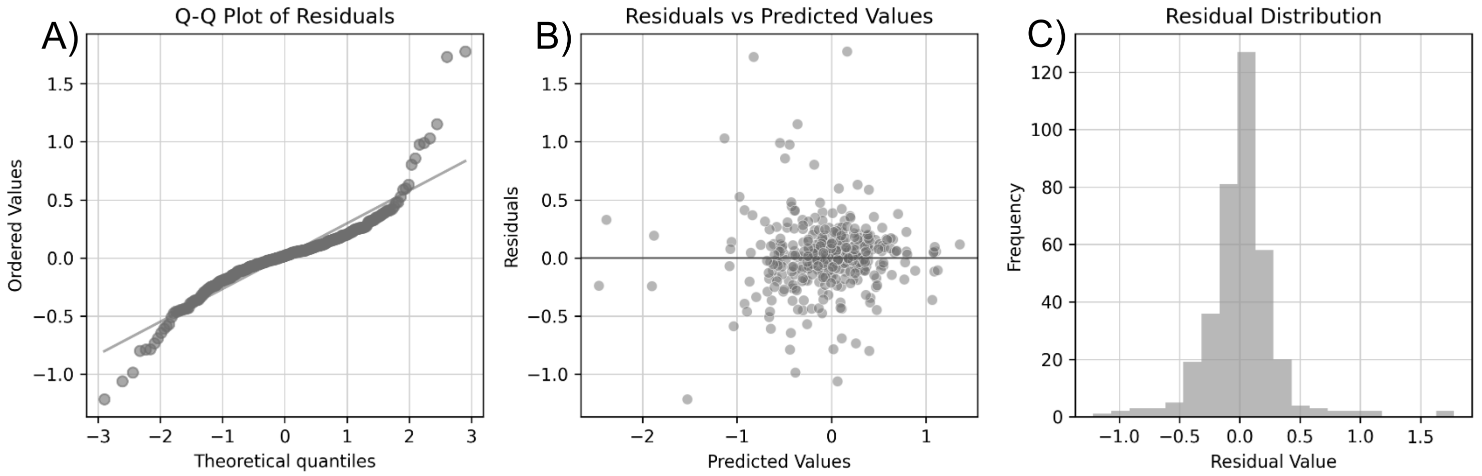


**Figure S9. Validation of the random forest model for predicting LFC.**

Diagnostic plots for the random forest model, which predicts plasmid log fold change (LFC) from assay design features. A) The Q-Q plot of residuals shows near-normality. B) The residuals vs. predicted values plot shows no clear pattern and consistent variance (homoscedasticity), indicating no systematic bias. C) The residual histogram is unimodal and centered at zero. Together, the plots confirm that the model's errors are well-behaved. Test Set Performance: $R^2 = 0.730$; MSE = 0.089; RMSE ≈ 0.299 LFC units.

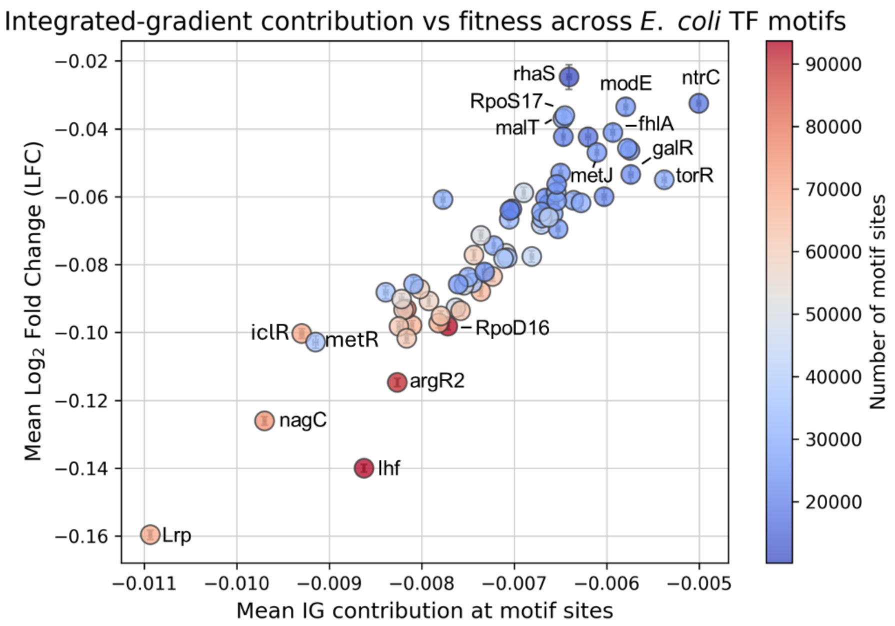


**Figure S10. Relationship Between Integrated Gradient (IG) scores and biological effect (LFC) for *E. coli* TFBS.**

Comparison between the transcription factor binding site's (TFBS) mean Integrated Gradient (IG) score with the mean $\log_2$ fold change (LFC) of the motifs across plasmids. Integrated Gradients quantify the direction and magnitude of each motif's contribution to the model's predicted fitness, with negative IG values indicating features that push predictions toward lower fitness, and positive IG values indicate features associated with higher predicted fitness. Each point represents a distinct *E. coli* TFBS with homologous sites found in the plasmid library. The IG and LFC values are averaged across all plasmids in which that motif occurs. LFC measurements were obtained in duplicate biological replicates. The colour scale reflects the total number of occurrences of each motif across the dataset, and error bars represent the standard error of the mean (SEM) for both IG and LFC.

### Supplementary Tables

**Table S1. Selected MTK plasmids for RNA sequencing analysis**

Twenty-four MTK were selected to span the space defined by predicted transcriptional load and observed fitness in DH5α (log fold-change, LFC), enabling analysis of constructs with high and low predicted transcription and contrasting fitness outcomes.

| MTK Plasmid | Description | Predicted TX | LFC in DH5a |
|---|---|---|---|
| MTK234_021 | U6-driven gRNA expression cassette | 596749.83 | 0.24 |
| MTK3_027 | Bicistronic ABI gene expression system | 2020663.28 | -0.19 |
| MTK3a_025 | Nuclear-localised deactivated Cas9 (dCas9) | 3241743.21 | -0.39 |
| MTK8b_004 | 3rd generation HIV-1 LTR promoter cassette | 2484221.69 | -3.93 |
| MTK3b_024 | Human NFATC2 transcription factor | 1416323.58 | -1.47 |
| MTK3_009 | Light-inducible VP16 transcriptional activator (PhyB fusion) | 2199578.11 | -4.12 |
| MTK3a_026 | Nuclear-localised synthetic dCas9 variant | 2006186.99 | -2.05 |
| MTK4a_021 | Nuclear-localised destabilised mRuby2 reporter | 865381.72 | -1.87 |
| MTK8b_005 | 5′ adeno-associated virus (AAV) ITR | 20963.44 | 0.30 |
| MTK5_011 | 5' Connector / UTR-like spacer (ConR8) | 82456.86 | -2.44 |
| MTK2_008 | Human pUbC | 749525.87 | 0.17 |
| MTK2_007 | Human pEF1α | 602783.59 | -0.34 |
| MTK3b_018 | Deactivated Cas9 (dCas9) | 2338745.81 | -0.33 |
| MTK3a_024 | Nuclear-targeted synthetic dCas9 | 3237289.50 | -0.39 |
| MTK3_033 | Gibberellin-inducible VPR transcriptional activator | 2087247.18 | -0.55 |
| MTK3b_030 | Dual-NLS deactivated Cas9 | 3256052.47 | -0.38 |
| MTK259 | pU6 | 467084.89 | 0.87 |
| MTK3b_019 | Synthetic deactivated Cas9 variant | 2006135.07 | -2.05 |
| MTK260 | PGKpA | 191057.31 | 0.26 |
| MTK3_037 | cAMP-responsive reporter module | 2703558.55 | -0.06 |
| MTK3a_013 | Monomeric green fluorescent protein (msGFP) | 294360.13 | -0.02 |
| MTK6_004 | Blue fluorescent protein and blasticidin resistance cassette | 1419647.50 | -1.33 |
| MTK4a_018 | Nuclear-localised cyan, fluorescent mTurquoise2 reporter | 1177012.13 | -0.41 |
| MTK5_010 | 5' Connector / UTR-like spacer (ConR7) | 87150.09 | -2.12 |
